# The influence of image masking on object representations during rapid serial visual presentation

**DOI:** 10.1101/515619

**Authors:** Amanda K. Robinson, Tijl Grootswagers, Thomas A. Carlson

## Abstract

Rapid image presentations combined with time-resolved multivariate analysis methods of EEG or MEG (rapid-MVPA) offer unique potential in assessing the temporal limitations of the human visual system. Recent work has shown that multiple visual objects presented sequentially can be simultaneously decoded from M/EEG recordings. Interestingly, object representations reached higher stages of processing for slower image presentation rates compared to fast rates. This fast rate attenuation is probably caused by forward and backward masking from the other images in the stream. Two factors that are likely to influence masking during rapid streams are stimulus duration and stimulus onset asynchrony (SOA). Here, we disentangle these effects by studying the emerging neural representation of visual objects using rapid-MVPA while independently manipulating stimulus duration and SOA. Our results show that longer SOAs enhance the decodability of neural representations, regardless of stimulus presentation duration, suggesting that subsequent images act as effective backward masks. In contrast, image duration does not appear to have a graded influence on object representations. Interestingly, however, decodability was improved when there was a gap between subsequent images, indicating that an abrupt onset or offset of an image enhances its representation. Our study yields insight into the dynamics of object processing in rapid streams, paving the way for future work using this promising approach.

## Introduction

The human brain processes rapidly changing visual input and can effortlessly extract abstract meaning when stimuli are presented in rapid sequences (Mack, Gauthier, Sadr, & Palmeri, 2008; Mack & Palmeri, 2011; Potter, Wyble, Hagmann, & McCourt, 2014; Thorpe, Fize, & Marlot, 1996; VanRullen & Thorpe, 2001). Recently, the temporal dynamics of the emerging representation of visual objects have been studied using fast presentation rates and multivariate analysis methods of electroencephalography (EEG) and magnetoencephalography (MEG) (Grootswagers, Robinson, & Carlson, 2019; Marti & Dehaene, 2017; Mohsenzadeh, Qin, Cichy, & Pantazis, 2018). Notably, multiple visual objects represented in different stages of the visual system can be decoded from the EEG signal at the same time (Grootswagers et al., 2019). Object representations persisted for longer when presented at slower presentation rates compared to faster rates (Grootswagers et al., 2019; Mohsenzadeh et al., 2018). Additionally, images presented at slower rates reached higher stages of processing, such that categorical abstraction of animacy was evident for images in 5Hz but not 20Hz rapid serial visual presentation (RSVP) sequences (Grootswagers et al., 2019). Previous studies have used time-resolved decoding to study perceptual versus conceptual representations (Linde-Domingo, Treder, Kerrén, & Wimber, 2019; Proklova, Kaiser, & Peelen, 2019; Ritchie & Op de Beeck, 2018; Teichmann, Grootswagers, Carlson, & Rich, 2018a, 2018b; Teichmann et al., 2019; Weaver, Fahrenfort, Belopolsky, & van Gaal, 2019). Changing presentation rates during RSVP offers the potential to explicitly target such different stages of processing. However, to do this we need to understand how exactly faster presentation rates influence image representations. The extended neural representations for slower versus faster presentation rates could be ascribed to the longer stimulus duration, or the longer stimulus onset asynchrony (SOA) of images in slower sequences.

Limitations in visual processing during RSVP is likely due to interference from processing multiple images in short succession. Decades of cognitive research have documented limitations in reporting targets during RSVP in phenomena such as the attentional blink (Broadbent & Broadbent, 1987; Raymond, Shapiro, & Arnell, 1992) and repetition blindness (Kanwisher, 1987). Such effects are typically studied to investigate high-level cognitive limitations rather than low-level visual processing interference (Raymond et al., 1992; Sergent, Baillet, & Dehaene, 2005). It is important to note, however, that target masking has a large effect on target detection during RSVP; for example, masking of the first and second targets increases the attentional blink deficit (Giesbrecht & Di Lollo, 1998; Nieuwenstein, Potter, & Theeuwes, 2009; Seiffert & Di Lollo, 1997). These findings suggest an important effect of low-level visual masking on higher-level processing during RSVP. In fast sequences, images are likely subject to forward masking by the previous image and backward masking by the next image in the sequence. Changing the image presentation rates has the effect of altering the timing of the masks. Backward and forward masking seem to have dissociable effects on perception, with one study showing maximal forward masking for 0ms gap between stimuli, and maximal backward masking at 30-90ms gap (Bachmann & Allik, 1976). EEG has shown that backward pattern masking influences processing after approximately 180ms, consistent with recurrent processing rather than feedforward processing deficits (Fahrenfort, Scholte, & Lamme, 2007). Understanding how masking affects the temporal dynamics of image processing during rapid-MVPA can yield important insights about the temporal limitations of the human visual system.

Studies of periodic visual evoked potentials also provide insights into the effect of image presentation rate on the extent of visual object processing. Faces presented at slower frequencies reach further stages of processing than those at faster rates, such that 15Hz presentations seemed limited to early visual processes, 6Hz showed increased occipitotemporal responses, and 3.75Hz included higher level cognitive effects and frontal responses (Collins, Robinson, & Behrmann, 2018). Retter et al., (2018) showed that SOA and image duration had dissociable effects on the periodic response. Images at 10Hz had larger evoked responses than those at 20Hz, but a 50% on-off image duty cycle (50ms duration, 100ms SOA) resulted in larger responses than 100% duty cycle with same SOA (100ms duration, 100ms SOA), a finding attributed to forward masking in the 100% duty cycle condition (Retter, Jiang, Webster, & Rossion, 2018). Taken together, it seems likely that SOA and image duration have separable influences on visual responses, but how these differentially influence the temporal dynamics of individual image processing remains to be seen.

Here, we investigate the effect of image masking on the temporal dynamics of image processing by studying the emerging neural representation of visual objects in fast visual streams while separately manipulating stimulus duration and SOA. These factors could be predicted to influence the temporal dynamics of individual image processing in a linear or non-linear fashion. Varying SOA, and thus the amount of time an image can be processed before another image (acting as a mask) appears, could linearly influence the duration of image processing if the length of processing is directly related to the amount of time dedicated to processing the uninterrupted images. Alternatively, there might be a limit on the number of items that can be held in the visual system at once. If SOA influences the dynamics of image processing depending on the stage of processing that is influenced by forward and backward masking, this would predict a non-linear increase in image processing. Our results show that a longer SOA enhances the decodability of the neural representations in a non-linear fashion, regardless of stimulus presentation duration. Our results also suggest that presenting stimuli with no gap between subsequent images (100% duty cycle) delays the processing of each image.

## Methods

Stimuli, data, and code are available at: https://doi.org/10.17605/OSF.IO/3RMJ9.

### Stimuli

We collected a stimulus set of 24 visual objects spanning 6 categories (Figure 1A). Stimuli were obtained from the free image hosting website www.pngimg.com. The top-level categories were animals and vehicles subdivided into 3 subcategories: birds, dogs, fish, boats, cars, and planes. Each of the subcategories consisted of 4 images each. These images allowed us to investigate visual representations for three different categorical levels: animacy (2 categories, animals/vehicles), object (6 categories, e.g., boats, birds) and image-level (24 images, e.g., yacht, duck). Images were presented using Psychtoolbox (Brainard, 1997; Kleiner et al., 2007; Pelli, 1997) in Matlab. Images were each shown foveally within a square at approximately 3 × 3 degrees of visual angle.

**Figure 1.**
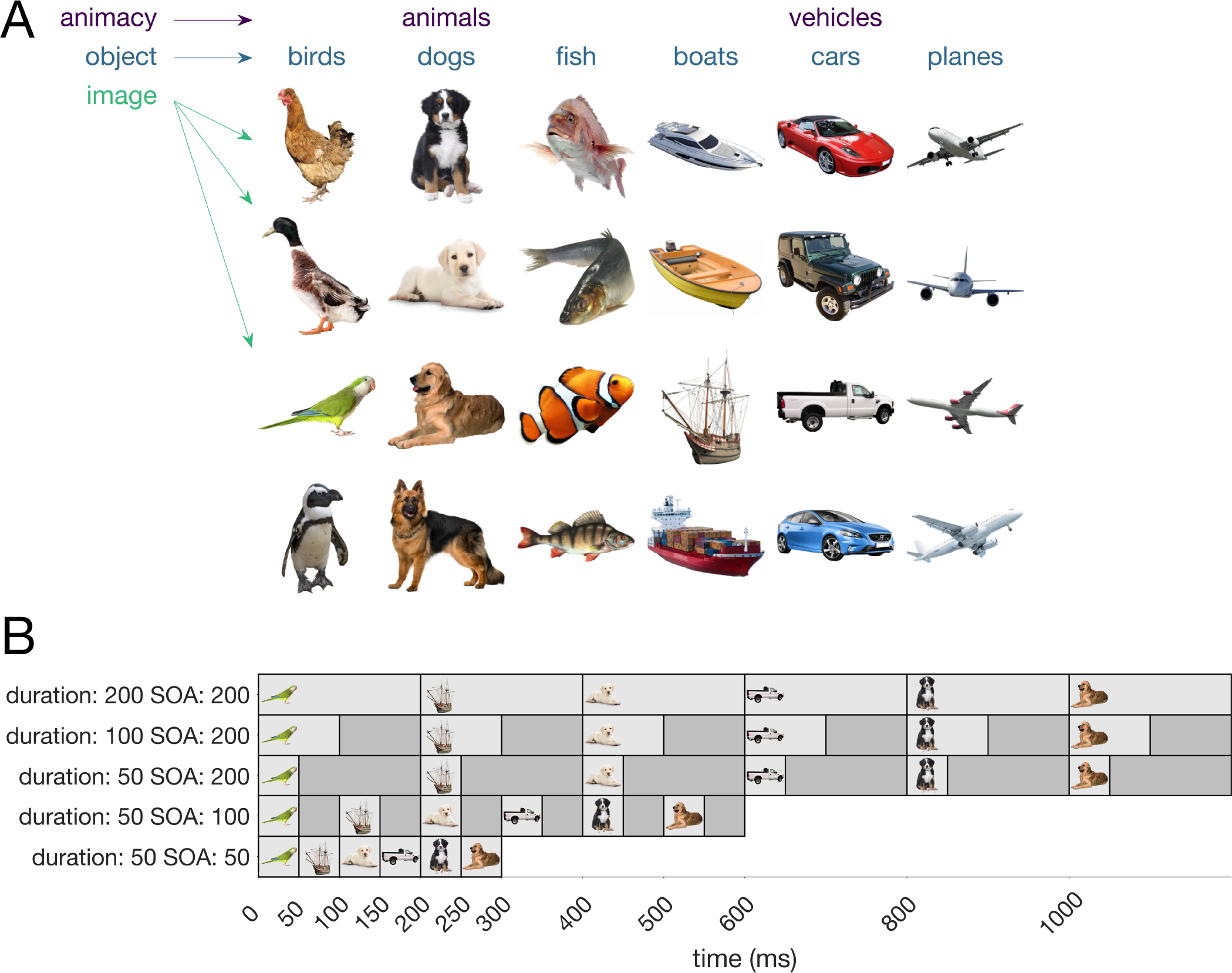
Stimuli and design. A) Experimental stimuli consisted of 24 images of objects organised at three different levels: animacy (animals, vehicles), object (6 categories e.g., birds, boats) and image (e.g., duck, chicken). B) Example time-lines illustrating the timing of the first six stimuli in a sequence in the different conditions. Images were presented in sequences with image durations of 200ms, 100ms and 50ms, and SOA of 200ms, 100ms and 50ms (5 conditions).

### Participants and experimental procedure

Participants were 20 adults recruited from the University of Sydney (12 female, 8 male; mean age: 25.75, age range 18-52 years) in return for payment or course credit. The study was approved by the University of Sydney ethics committee and informed consent was obtained from all participants. Participants viewed 200 sequences of objects. Each sequence consisted of the 24 stimuli in a random order. To ensure all images were equally masked by other images, the sequences were padded with 12 stimuli on both ends, which were excluded from the decoding analysis. The 12 padding stimuli consisted of the same sequence in reverse order, with mirrored versions of the images. The mirror-reverse padding ensured a minimum of 12 images between two repeats of the same image within a sequence and that each of the 24 experimental images was presented twice per sequence. To keep participants engaged, at the end of each sequence, after a 1000ms blank screen, a random image from the stimulus set was presented for 100ms and participants categorised this stimulus as animal or vehicle using a left or right button press (response mappings were alternated between participants). The presentation rates of the sequences were chosen from one of five conditions, which were randomized throughout the study (40 sequences per condition). In conditions 1-3, the presentation duration varied (200ms, 100ms, and 50ms) while keeping the SOA at 200ms. In conditions 3-5, the SOA varied (200ms, 100ms, and 50ms) while keeping the presentation duration at 50ms (Figure 1B). This set-up allowed us to use condition 3 as anchor point to compare the effects between varying SOA and duration. In total, participants viewed 9600 presentations, consisting of 80 presentations for each of the 24 images and for the 5 duration/SOA conditions.

### EEG recordings and preprocessing

EEG data were continuously recorded from 64 electrodes arranged in the international 10–10 system (Oostenveld & Praamstra, 2001) using a BrainVision ActiChamp system, digitized at a 1000-Hz sample rate. Scalp electrodes were referenced to Cz during recording. EEGlab (Delorme & Makeig, 2004) was used to pre-process the data offline, where data were filtered using a Hamming windowed sinc FIR filter with highpass of 0.1Hz and lowpass of 100Hz. Data were then downsampled to 250Hz and epochs were created for each stimulus presentation ranging from [−100 to 1000ms] relative to stimulus onset. The same epoch length was used for all conditions. No further preprocessing steps were applied.

### Decoding analysis

An MVPA time-series decoding pipeline (Grootswagers, Wardle, & Carlson, 2017; Oosterhof, Connolly, & Haxby, 2016) was applied to each stimulus presentation epoch in the sequence to investigate object representations in fast sequences. Voltages for all 63 EEG electrodes were used as features for the decoding analyses, which was performed for each time point separately. Linear discriminant analysis classifiers were trained using an image by sequence cross-validation procedure (Grootswagers et al., 2019) to distinguish between all pairwise groupings within the categorical levels (animacy, object). This entailed holding out one image from each category in one sequence as test data and training the classifier on the remaining images from the remaining sequences. For pairwise decoding of the non-categorical image-level, we used a leave-one-sequence-out cross-validation procedure. It is important to note that the randomised image presentation order within each sequence ensured that for any given image, the preceding and following images were not informative to the classifier.

The decoding analyses were performed separately for the five duration/SOA conditions. For each condition, this resulted in three decoding accuracies over time (for animacy, object, and image). At each time point, these accuracies were compared against chance (50%) and compared to each other. All steps in the decoding analysis were implemented in CoSMoMVPA (Oosterhof et al., 2016).

### Statistical inference

We used Bayes factors (Dienes, 2011, 2016; Jeffreys, 1961; Kass & Raftery, 1995; Rouder, Speckman, Sun, Morey, & Iverson, 2009; Wagenmakers, 2007) to determine the evidence for the null and alternative hypotheses. For the alternative hypothesis of above-chance decoding, a uniform prior was used ranging from the maximum value observed during the baseline (before stimulus onset) up to 1 (i.e., 100% decoding). For testing the hypothesis of a difference between decoding accuracies, a uniform prior was set ranging from the maximum absolute difference between decoding accuracies observed during the baseline up to 0.5 (50%). We then calculated the Bayes factor (BF), which is the probability of the data under the alternative hypothesis relative to the null hypothesis. We thresholded BF>6 as strong evidence for the alternative hypothesis, and BF<1/6 as strong evidence in favour of the null hypothesis (Jeffreys, 1998; Kass & Raftery, 1995; Wetzels et al., 2011). BF that lie between those values indicate insufficient evidence for either hypothesis.

To determine onset, offset, and peak time signatures, we defined onset as the second time point where the Bayes factor exceeded 6 and offset as the second-to-last time point where the Bayes factor exceeded 6. Peak decoding time was defined as the latency at which the maximum decoding accuracy was observed in the entire time window. We obtained bootstrap distributions of these latency measures by sampling from the participants with replacement 1000 times and recomputing the abovementioned statistics.

### Exploratory channel searchlight analysis

To explore the spatial source underlying our decoding results, we performed a time-by-channel searchlight analysis. For every channel, a decoding analysis was performed using the data from a local five-channel cluster that included the channel’s four closest neighbours (Oostenveld, Fries, Maris, & Schoffelen, 2010; Oosterhof et al., 2016). This was repeated for each categorical contrast yielding a time-by-channel map of decoding accuracies. Instead of obtaining pairwise decoding accuracies, we computed the multiclass decoding accuracies to save computation time.

## Results

We examined the temporal dynamics of object processing using rapid-MVPA with sequences of varying image duration and SOA. During the experiment, participants reported whether an image presented after each sequence was an animal or a vehicle. Behavioural performance was high for discrimination of animal (*M* = 97.10%, *SD* = 5.23%) and vehicle (*M* = 98.10%, *SD* = 3.01%) stimuli.

To investigate the temporal dynamics of object processing, we decoded the objects at three levels of categorical abstraction: animacy-level (animal versus vehicle), object-level (birds, dogs, fish, boats, planes, cars), and image-level (24 images; 4 per object). The decoding analyses were performed separately for each SOA and image duration condition. Figure 2 shows the temporal dynamics of all categorical representations varied by SOA and image duration. For the effect of duration (left columns), all durations followed a similar decoding trajectory, but classification was poorer in general for the longest duration, which also happened to be the 100% duty-cycle condition (200ms SOA, 200ms duration). For animacy and object decoding, the first peak (~100ms) was similar across the conditions, but the 200ms duration was lower than both the 50ms and 100ms conditions from 150-200ms, suggesting poorer categorical abstraction for this condition. Additionally, for the individual image decoding analysis, the onset of decoding appeared to be delayed for the 200ms duration relative to the 50ms and 100ms durations.

**Figure 2.**
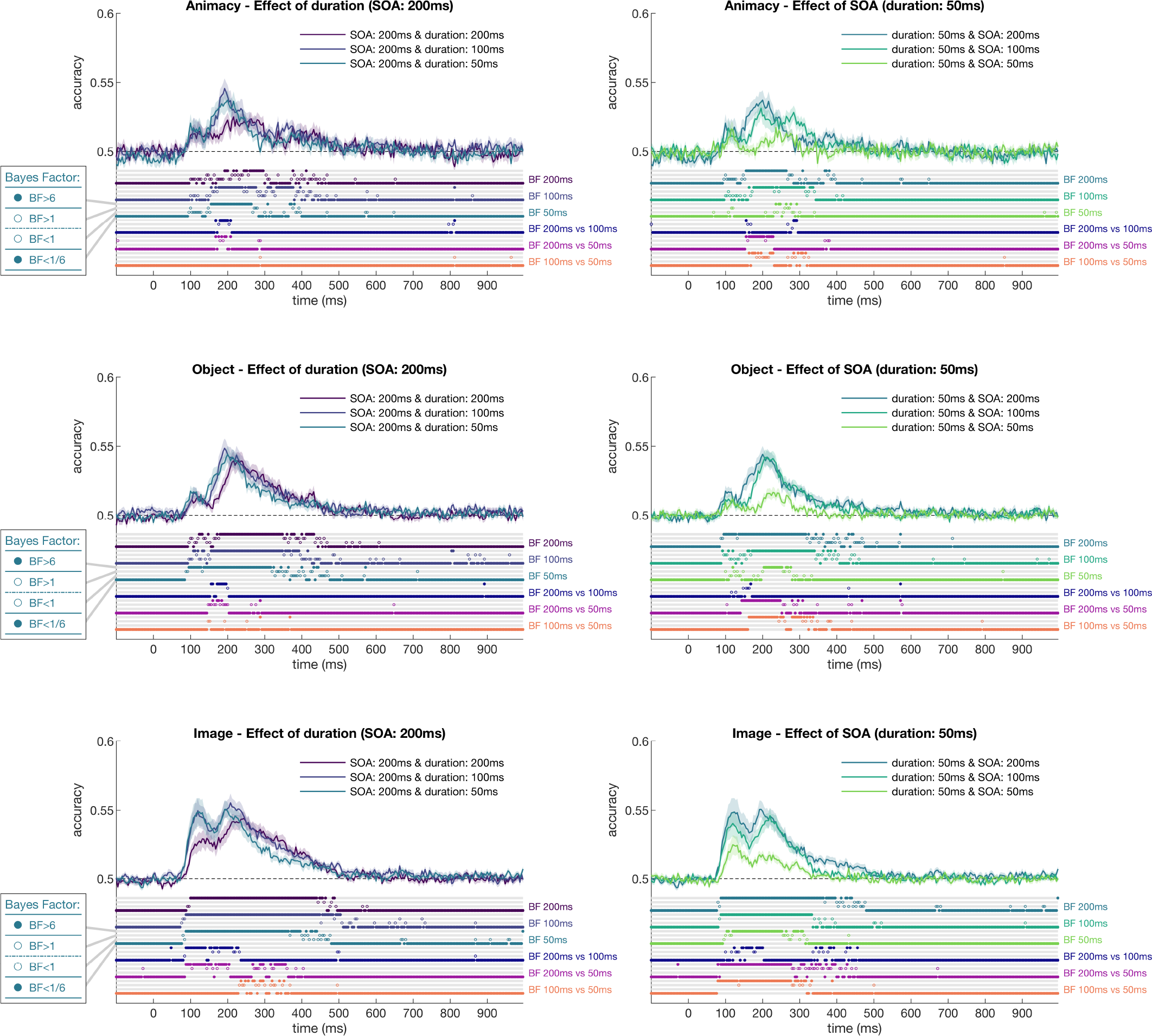
The effects of duration (left column) and stimulus onset asynchrony (right column) on decoding accuracy at three categorical levels (animacy, object, and image). Dots above the x-axis show the thresholded Bayes factors for every time point as an open or closed circle in one of four locations (see insets), reflecting the amount of evidence for the null or alternative hypothesis. The top three groups of Bayes factors show the evidence for above-chance decoding, and the bottom three show Bayes factors for differences between decoding accuracies.

The right columns of Figure 2 show that for a given image duration (50ms), increasing SOA led to greater neural decoding for all categorical levels. For animacy and object decoding, the initial peak (~120ms) did not differ by SOA, but the larger second peak (~200ms) showed graded responses depending on SOA. At this peak, there was a small but reliable increase in decoding for the 200ms SOA relative to the 100ms, and these were both substantially higher than the 50ms SOA. Again, for the image decoding the 100% duty-cycle condition (50ms SOA/50ms duration) appeared delayed and had poorer decoding relative to the other conditions. Furthermore, the 200ms SOA had greater decoding than the 100ms SOA condition between 100 and 200ms. Overall, these results imply that longer SOA led to stronger image representations.

To further assess the effect of image duration and SOA on object decoding, we analysed the timing of the decoding window (onset to offset of above-chance decoding) and the latency of peak decoding. Figure 3 shows that the medium duration condition (duration 100/SOA 200) had the longest decoding window for all decoding contrasts. In contrast, the shortest duration and SOA condition (duration 50/SOA 50) had delayed onsets and the shortest decoding window. The peak latency results revealed that for animacy and object, the 100% duty-cycle conditions (200/200 and 50/50) had the latest peaks. For the image decoding, again the 200/200 condition had the latest peak, whereas the 50/50 condition had a much earlier peak. This seems to be due to limited ongoing processing in the 50/50 condition such that peak decoding was for the first decoding peak whereas the other conditions had larger second peaks. In essence, it seems that the 50/50 peak was actually centred on a different process than the peaks for the other conditions. Nevertheless, the *onset* of processing appears to be consistently delayed for the 100% duty-cycle conditions.

**Figure 3.**
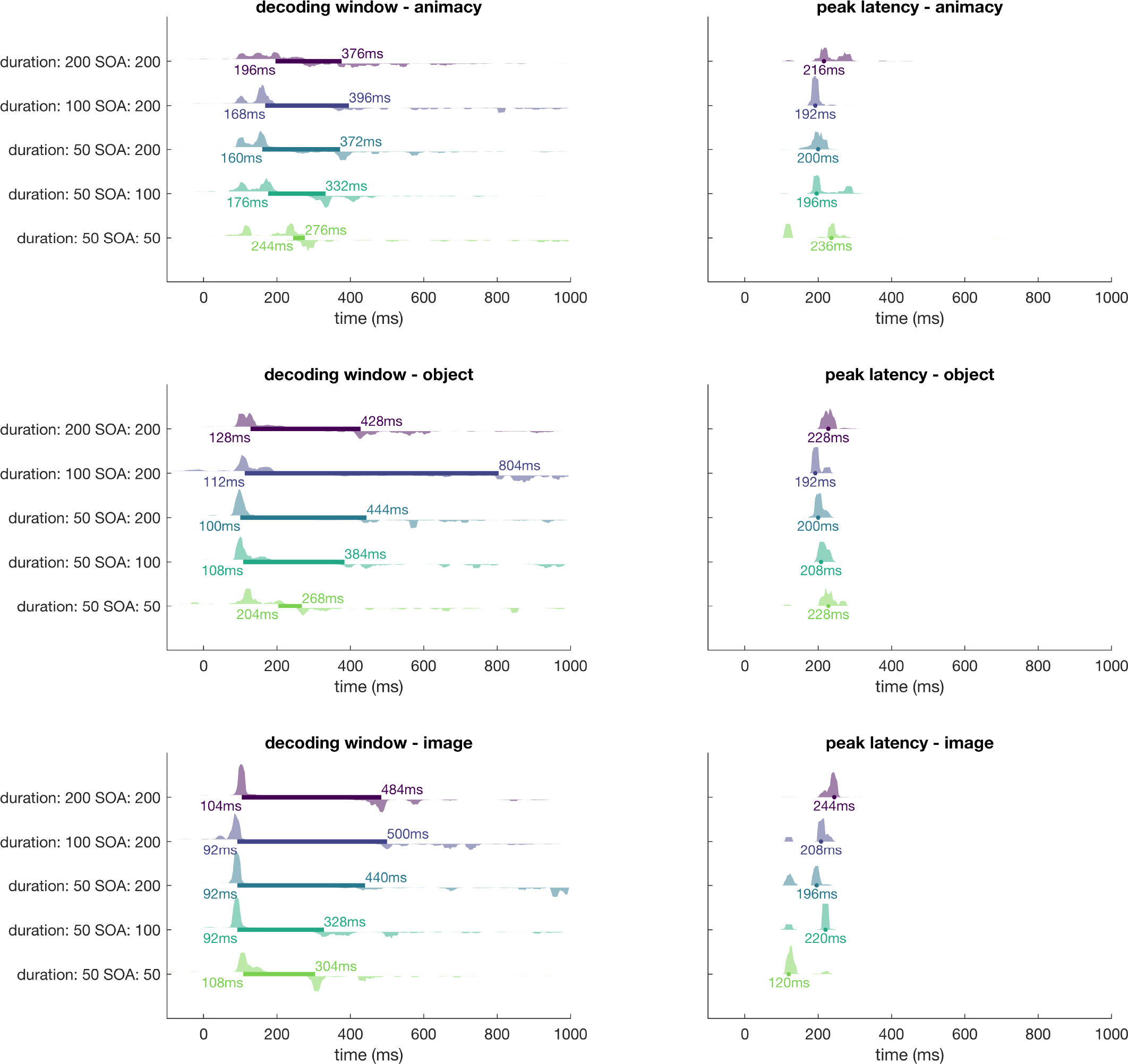
Onset, offset, and peak latencies for each condition. Onset was defined as the second time point where the Bayes factor exceeded 6 and offset as the second-to-last time point where the Bayes factor exceeded 6. The three rows show the result for the three categorical levels (animacy, object, and image). Left columns: for each condition (y-axis), onset and offset are marked by a filled horizontal bar and are annotated at their respective time points. Shaded areas show the onset (above the filled bar) and offset (below the filled bar) latency distributions calculated by bootstrapping participants with replacement 1000 times and recomputing the statistics. Right columns: Peak latency (time point of peak decoding) for each condition (y-axis, same order as the left column). Shaded areas show the peak latency distribution calculated by bootstrapping participants with replacement 1000 times.

To explore the spatial origins of the decodable information, we performed a time-by-channel searchlight. Figure 4 shows the averaged multiclass decoding accuracies of this searchlight for image decoding at the time windows that correspond to the two peaks observed in Figure 2 for all five conditions. For all categorical levels and timepoints, see https://doi.org/10.17605/OSF.IO/3RMJ9. During the first peak (100-150ms), the signal was mainly located in central occipital sensors, and during the second peak (175-225ms) in occipitotemporal sensors. The second window suggested that stronger accuracies were lateralised towards the right hemisphere.

**Figure 4.**
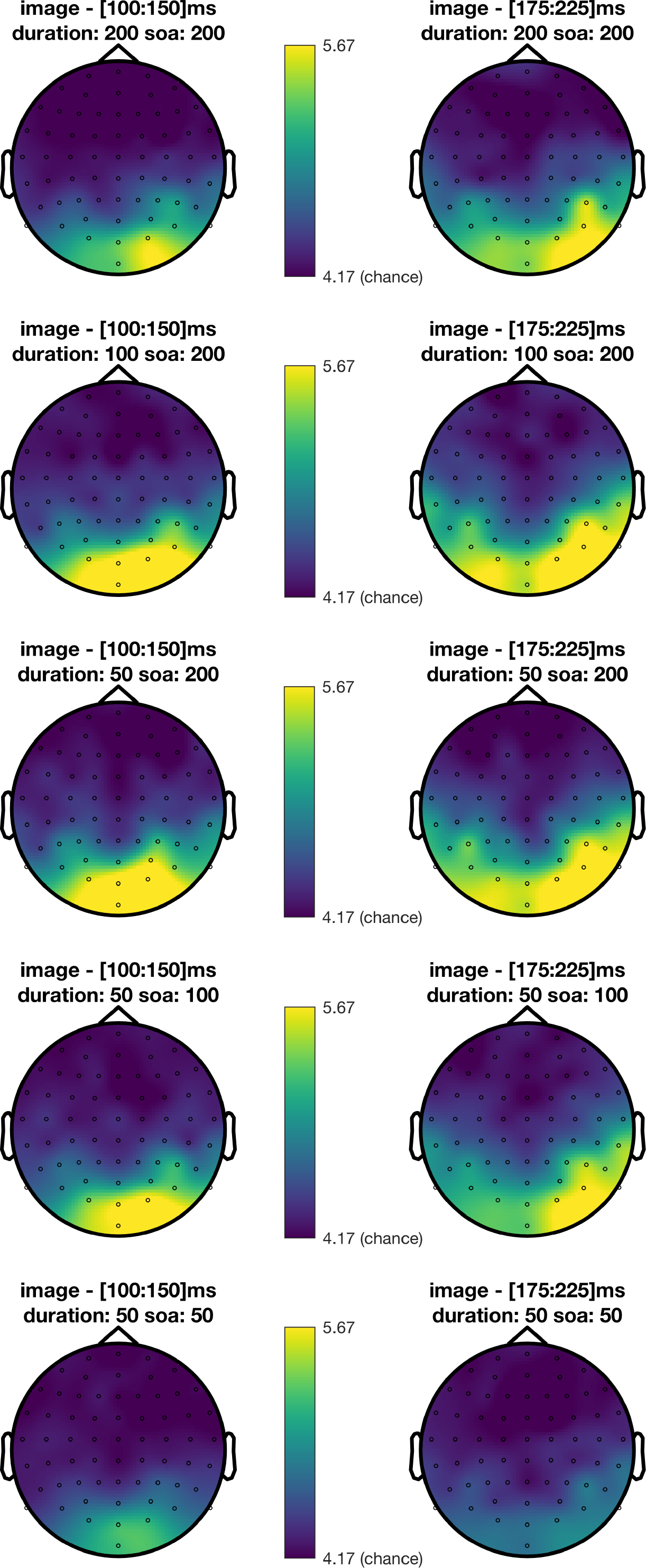
Exploratory channel-searchlight analysis. Plots show average multiclass image decoding accuracies as a topographic distribution over the head. Rows show all five presentation conditions. Columns show two time windows that correspond to the peak decoding time points in Figure 2. Chance-level is 4.17% (i.e., 1/24).

## Discussion

In this study, we disentangled the effects of duration and stimulus onset asynchrony (SOA) on decoding performance in rapid visual processing streams. Our results showed that shorter SOAs systematically reduced the duration of above-chance decoding, as well as the peak decoding accuracy, consistent with masking at earlier stages of visual processing. In comparison, there were no graded effects of presentation duration on decoding accuracies. Our results also suggest that presenting stimuli without a gap (100% duty cycle) leads to delays in visual processing.

Previous work found that fast presentation rates limits visual processing relative to slower presentation rates (Grootswagers et al., 2019). It was however unclear whether this difference was due to shorter stimulus duration or shorter SOA. The results of our study show that stimulus duration and SOA have separable effects on stimulus processing, with the most pronounced effect being that longer SOAs enhance decodability of stimuli relative to shorter SOAs. These findings are consistent with recent work that investigated the effect of duration and SOA on face response amplitudes in a fast periodic visual stimulation paradigm (Retter et al., 2018). Single unit recordings in temporal cortex of macaques have revealed a similar effect; neural responses to monkey faces in RSVP are stronger and last longer for slower presentation rates (Keysers, Xiao, Földiák, & Perrett, 2001, 2005). Interestingly, Keysers et al. (2001) found that the duration of image discrimination coincided with the SOA length plus 60 ms, an effect attributed to neural competition with other images in the sequence (Keysers & Perrett, 2002). Although we found that longer SOA led to longer neural decoding, there was no clear linear relationship between the SOA and length of decoding. Our decoding results utilise whole brain responses, however, which might be one reason for this difference; It could be, for instance, that non-linear interactions between early and late visual cortical regions overshadow or obscure linear effects within any one region as measured with EEG. The current results suggest that SOA influences the degree of masking from subsequent images depending on the stage of processing that is disrupted.

Our findings that SOA influences object representations in RSVP is consistent with a backward masking account, such that presentation of every new image impairs processing of the previous image. The mechanism for this masking could be explained by conceptual masking, neural competition, or interruption. Conceptual masking is a high-level type of masking observed when the critical image and the mask both activate high level concepts that compete for resources (Intraub, 1984; Potter, 1976). It seems unlikely that our observed effects could be described by conceptual masking because we see a graded effect of SOA on the amplitude and duration of information coding, rather than a common stage of processing that is disrupted. As an alternative, the interruption and neural competition accounts of masking imply that capacity limited processes within the visual hierarchy mean more than one image cannot be fully processed within a short period of time. Our results are consistent with capacity limits within the human brain, however it is important to note that our results suggest that multiple object representations can co-exist in the brain simultaneously, because decoding duration outlasted many subsequent image presentations. What seems likely is that competition and/or interruption can occur within high-order brain regions (Keysers & Perrett, 2002) but at any one time, different object representations can be present at different stages of processing (and potentially within different brain regions). Because of the whole brain approach of EEG decoding, we cannot determine whether competition or interruption are more likely processing for the masking effects observed in the current study, but future work could investigate this using a more fine-grained approach with many more combinations of duration and SOA.

In contrast to the effects of SOA, we did not observe a notable effect of stimulus duration on object representations during RSVP. This is consistent with neural persistence, such that visual information continues to be processed even when it is no longer visible (Duysens, Orban, Cremieux, & Maes, 1985). Behaviourally, research has found that images presented for a short duration with a large gap are remembered almost as well as images presented for long durations (Potter, 2012; Potter, Staub, & O’connor, 2004). Our neural results add to such findings by showing that objects presented for short durations followed by a blank gap (e.g., 100 duration/200 SOA vs 50 duration/200 SOA) do not appear to be processed differently. The notable exception to this effect was for sequences with no gap between successive images. Specifically, neural responses were delayed when images were presented back-to-back (100% duty cycle). The most likely explanation for this is forward masking, such that processing of an image impaired processing of the next image, which is also supported by behavioural results (Bachmann & Allik, 1976). Furthermore, in EEG, smaller periodic signals have been found with 100% duty cycles than 50% duty cycles (Retter et al., 2018). Our results suggest that forward masking impairs perception by delaying neural responses to the next image. Taken together, the findings of this study elucidate some of the complex neural mechanisms operating during RSVP which include neural persistence, forward masking and backward masking, each of which appear to exhibit differential effects on object representations.

Our results can also be used to guide future visual object representation studies that employ a rapid-MVPA design. For example, using a 5Hz rate, Grootswagers et al. (2019) obtained 40 epochs for 200 stimuli (8000 epochs in total) in a 40-minute session. Here, we did not observe strong differences between 5Hz and 10Hz presentation rates (200ms and 100ms SOA). Thus, a 10Hz 50% duty cycle presentation paradigm seems to provide a sensitive measure of object decoding accuracy. Notably, this is also a typical frequency used in RSVP paradigms to study target selection processes, which are postulated to involve alpha oscillatory activity (Janson, De Vos, Thorne, & Kranczioch, 2014; Zauner et al., 2012). At 10Hz, a 30-minute EEG recording session (excluding breaks) yields 18000 epochs, which has unprecedented potential for studying a large number of different conditions and/or stimuli. It also suggests that it is possible to obtain enough epochs for a small number of conditions in a very short (<5-minute) EEG session. This opens up exciting new possibilities to study special populations for whom long experiments often pose significant difficulties, such as children and patients.

This study showed that analysing the neural signatures of all images in RSVP streams can yield insight into the mechanisms underlying visual masking. Without the need of a separate noise mask, a substantial number of presentations or conditions can be tested using rapid-MVPA, increasing the power of such experiments. By shortening the SOA, the masking affected earlier stages of processing, which has significant potential for studying hierarchical processing systems, such as vision (see also McKeeff, Remus, & Tong, 2007). For example, future work could apply rapid-MVPA and varying SOAs to stimulus sets that vary on orthogonal features that are expected to occur at different stages in the processing streams, such as colour and shape.

## Acknowledgements

The authors thank Ayeesha Dadamia for assisting with data collection, and Denise Moerel for helpful comments on a draft of the manuscript. This work was supported by an Australian Research Council Future Fellowship (FT120100816) and an Australian Research Council Discovery project (DP160101300) awarded to T.A.C. The authors acknowledge the University of Sydney HPC service for providing High Performance Computing resources. The authors declare no competing financial interests.

